# Exogenous calcium ions enhance patulin adsorption capability of *Saccharomyces cerevisiae*

**DOI:** 10.1101/391573

**Authors:** Ying Luo, Xiaojiao Liu, Yanqing Han, Jianke Li

## Abstract

Patulin contamination is a severe issue that restricts the development of the global fruit processing industry. Yeast adsorbs patulin more effectively than other microbial adsorbents, and this adsorption process mainly depends on the function of the cell wall. Additionally, exogenous calcium ions aid in yeast cell wall formation according to reports. Therefore, in the present study, the effect of exogenous calcium concentrations on the cell wall structure and the patulin adsorption capability was studied. We showed that the ability of the yeast to adsorb patulin was strengthened with an increase in exogenous calcium concentrations between 1×10^-4^ - 1×10^-2^ mol/L. Moreover, yeast cell wall thickness, β-1,3-glucan content and the activities of the key catalytic enzymes β-1,3-glucanase and β-1,3-glycosyl transferase were all increased within this range. The results indicated that exogenous calcium activates key enzymes and that these enzymes are crucial for cell wall network formation and patulin adsorption capability.

**Importance:** The present work illuminates that the exogenous calcium ions could determine the insoluble network structure by regulating key enzyme activities under certain concentrations, thus indirectly influencing the yeast cell patulin adsorption capability. It could enhance patulin adsorption capability of yeast walls and successfully apply to fruit juice industry.

## 1. Introduction

Patulin is a kind of secondary metabolite produced by a wide range of fungi during their growth on rotting fruit. The presence of patulin in fruit and vegetable products, especially in apple products, has become a severe problem for food safety. It has been reported that approximately 50% of the analyzed samples showed relatively high detectable patulin levels in apple juice worldwide ^[1]^. This mycotoxin causes acute and chronic damage in animal studies and in vitro experiments ^[2, 3]^. Patulin was classified as a category 3 toxin by the International Agency for Research on Cancer ^[4]^, and the European Union (EU) has recommended that the maximum patulin detection levels are 50 μg/kg for fruit juices and 10 μg/kg for infant products ^[5]^.

To ensure fruit product safety, numerous approaches, including physical, chemical, and biological methods, have been developed to eliminate patulin occurrence. Biological adsorption has recently been considered the most effective strategy in the food industry. Yeast adsorption is considered to have a dominant role compared to other microorganisms due to its unique advantages, such as easier cultivation, lower cost, and lack of hazards ^[6]^. Most yeast species, such as *Saccharomyces cerevisiae, Candida* spp., *Pichia* spp., and *Rhodotorula* spp., can adsorb patulin and other mycotoxins ^[6, 7]^. The yeast cell wall allows cells to adsorb a range of compounds from the environment, and it was reported to be the major component for patulin adsorption. In addition, we could state that the β-glucans that make up the cell wall play an important role in patulin adsorption ^[8]^. Yiannikouris and others indicated that yeast strains possess a larger number of β-glucans and a greater amount of chitin and thus were able to adsorb larger amounts of mycotoxins ^[9]^. Furthermore, the key factors in the yeast cell wall β-glucan and mycotoxin adsorption processes were identified as van der Waals and hydrophobic interactions ^[10]^. Based on these facts, the interactions between the β-glucans and patulin are more of an adsorption type with physical interactions, and the three-dimensional network structure of β-glucans has an important role in the adsorption.

The yeast cell wall three-dimensional network structure mainly consists of β-1,3- and β-1,6-glucan chains linked to chitin and mannoproteins ^[11]^. The network was identified as having uracil diphosphate-glucose (UDP-glucose) as its unique precursor substance, and β-1,3-glucan soluble chains were then biosynthesized utilizing β-1,3-glucanase. Next, β-1,3-glucan insoluble chains were generated by cross-linking with chitin catalyzed by β-1,3-glycosyl transferase ^[12]^ (Figure 1). Some research indicated that adding an appropriate amount of calcium ions during the yeast growth process promoted β-1,3-glucan formation and the activity of β-1,3-glucanase, thus predicting that it may be associated with calcium signaling pathways ^[13]^. Since calcium ions play a critical role as intracellular messengers in eukaryotic cells, they could allow the activation of the target protein calcineurin (CaN), which carries out multiple functions, including cell wall formation ^[14–16]^. Nevertheless, the manner in which exogenous calcium influences the patulin adsorption capability of the yeast cell wall and the specific relationship between them have rarely been reported.

**Figure 1.**
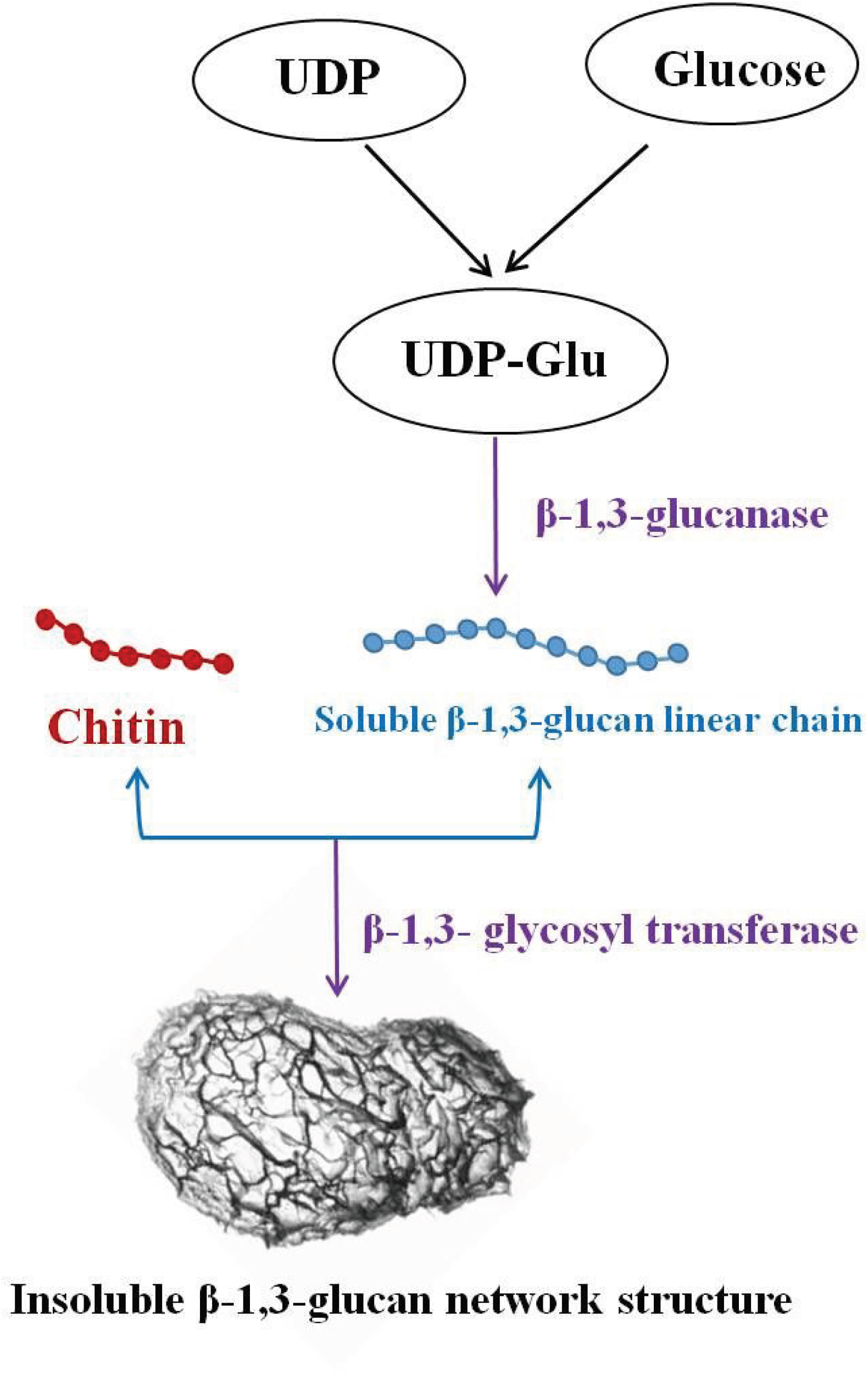
Yeast cell wall network structure biosynthesis and key enzymes involvement.

To explore the relationship between the exogenous calcium concentration and patulin adsorption capability of the yeast, this study aims to 1) verify the effect of exogenous calcium concentrations on the yeast cell wall yield and cell wall thickness, 2) illuminate the role of exogenous calcium concentrations on yeast cell wall β-1,3-glucan content and β-1,3-glucanase activity, and 3) analyze the patulin adsorption improvement mechanism.

## 2. Materials and methods

### 2.1 Materials and reagents

Standards of patulin, standard β-glucan, and calcium chloride were all purchased from Sigma-Aldrich (St. Louis, MO, USA). *Saccharomyces cerevisiae* ATCC 18824 was purchased from the American Type Culture Collection. Fluo 3-AM, and Pluronic F127 were obtained from Solarbio (Beijing, China). Other chemicals used in the experiments were all obtained from a local chemical reagent company. All chemical reagents were of analytical grade, and the solutions were prepared with deionized water.

### 2.2 Different concentrations of exogenous calcium and the preparation of cultivated yeast cells preparation

Yeast cells were cultivated in yeast extract peptone dextrose medium (YPD culture medium: glucose 2%, peptone 2% and yeast extract powder 1%) at 120 rpm, 30°C for 24 h. After cultivation was activated, yeast cells (5% inoculum size) were inoculated into calciferous YPD culture media with exogenous calcium concentrations ranging from 1×10^-4^ mol/L to 1 mol/L (120 rpm, 30°C for 24 h). After calcium cultivation, the cells were collected by centrifugation and washed twice with sterilized water. To analyze the yeast cell biomass (g/L), the cells were collected from 1000 mL of cell suspension and weighed after being freeze-dried to a constant weight.

### 2.3 Intracellular calcium concentration determination

To determine the intracellular calcium concentration, yeast protoplasts cultivated in different amounts of calcium were incubated at 37°C for 30 min with the addition of the calcium fluorescence probe Fluo 3-AM. Cells loaded with Fluo 3-AM then adhered to microscope slides using polylysine. Laser confocal fluorescence microscopy was used to determine the concentration of intracellular calcium. The excitation and emission wavelengths of Fluo 3-AM were 488 nm and 525 nm, respectively.

### 2.4 Cell wall morphology and thickness analysis

The cell wall thickness was determined using transmission electron microscopy (TEM) (JEOL-1230; JEOL Ltd., Japan). Different calcium-cultivated yeast cells were used to prepare the specimens for TEM. Thirty cells were randomly selected from five different fields of view. For each cell, four different points were measured. The cell wall thickness statistics were obtained using a frequency histogram.

### 2.5 Cell wall network structure components analysis

Different calcium-cultivated yeast cells were disrupted using an ultrasonic cell disruption system (Scientz-IID, Ningbo Xinzhi Biotechnology Co., Ltd.). For 1.3-β-glucan and 1,6-β-glucan extraction and purification, the cell wall fractions were extracted with NaOH at 75°C. The alkali-insoluble and alkali-soluble glucans were 1.3-β-glucan and 1,6-β-glucan, respectively ^[17]^. A Dionex Bio-LC system (ICS2500, USA) coupled with an ED 50 electrochemical detector was used to quantitatively analyze the cell wall carbohydrates. Deionized water: 0.5 mol/L NaOH (3.5: 96.5; v/v) was used as the isocratic mobile phase with a flow rate of 1 mL/min at room temperature. The chitin content was determined using an enzymatic method as described in other reports ^[18]^.

### 2.6 Extraction and activity determination of β-glucanase and β-1,3-glycosyl transferase

A spectrophotometric method was used to detect β-glucanase activity. Resuspended yeast cells grown in differing concentrations of calcium had their cell walls disrupted with glass beads. The samples diluted with phosphate buffer were mixed with lichenan at 50°C for 10 min, and 3,5-dinitrosalicylic acid was then added and boiled for 5 min. Spectrophotometry was used to measure the β-glucanase activities at 520 nm ultraviolet wavelength after the samples had cooled. A specific assay kit was prepared to detect β-1,3-glycosyl transferase activity after cell walls disrupted with glass beads.

### 2.7 Patulin adsorption and analysis in aqueous solution and apple juice

Different concentrations of calcium-cultivated yeast cells were suspended in 200 μg/L aqueous solution (1 mL) and patulin-contaminated apple juice (10 mL) with 10^6^/mL and 100 mg yeast cell addition, respectively. The cells were incubated for 10 h (150 rpm at room temperature) in a shaker incubator. The control sample lacked added cells. Three replications were prepared for each sample, and the independent experiments were performed three times. After 10 h, the cells were separated by centrifugation at 3600×g for 5 min, and the supernatants were then collected to extract and detect patulin ^[19]^. Patulin was analyzed by high-performance liquid chromatography (HPLC), separated by a C18 reversed-phase column, and detected using HPLC connected with a UV absorbance detector set at 276 nm. An acetonitrile: water solution (10: 90) was used as the isocratic mobile phase with a flow rate of 1 mL/min at 30°C, and the elution time was 15 min for each sample ^[20]^. The patulin adsorption efficiency (R%) was calculated using the following equation:

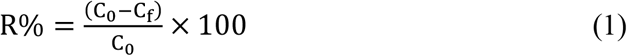

where *C_0_* and *C_f_* are the initial and final concentrations of patulin (mg/L), respectively.

### 2.8 Statistical analysis

The experiments were generally performed in triplicate, and the data are presented as the mean ± standard deviation. All data were subjected to one-way ANOVA using the Statistical Analysis System (SAS Inst., Cary, N.C., U.S.A.). The data were considered statistically significant when *p* < 0.05.

## 3. Results and discussion

### 3.1 The effect of exogenous calcium concentrations on yeast cells and cell wall biomass

The calcium ion is an essential element that serves as an intracellular messenger in yeast cells ^[21]^. Adding exogenous calcium during yeast growth contributes to the activation of the calcium pathway, thus regulating cell growth and cell wall synthesis ^[22]^. Changes in yeast cells and their cell wall biomass cultivated with different concentrations of exogenous calcium are compared in Figure 2. Yeast cell and cell wall biomass were positively correlated with exogenous calcium concentrations at a range from 1×10^-4^ mol/L to 1×10^-2^ mol/L, with the highest biomass of 3.93 g/L for the yeast cells and 0.38 g/L for the cell walls at the calcium concentration of 1×10^-2^ mol/L. With calcium concentrations continuing to increase, the yeast cell biomass subsequently decreased slightly at a calcium concentration of 1×10^-1^ mol/L and then decreased rapidly when the calcium concentration increased to 1 mol/L with a biomass of only 1.34 g/L. However, the cell wall biomass suddenly decreased from 1×10^-1^ to 1 mol/L, finally dropping to 0.24 g/L. The results indicated that exogenous calcium could enhance yeast and yeast cell wall growth in a certain extent and that the most effective concentration was 1×10^-2^ mol/L.

**Figure 2.**
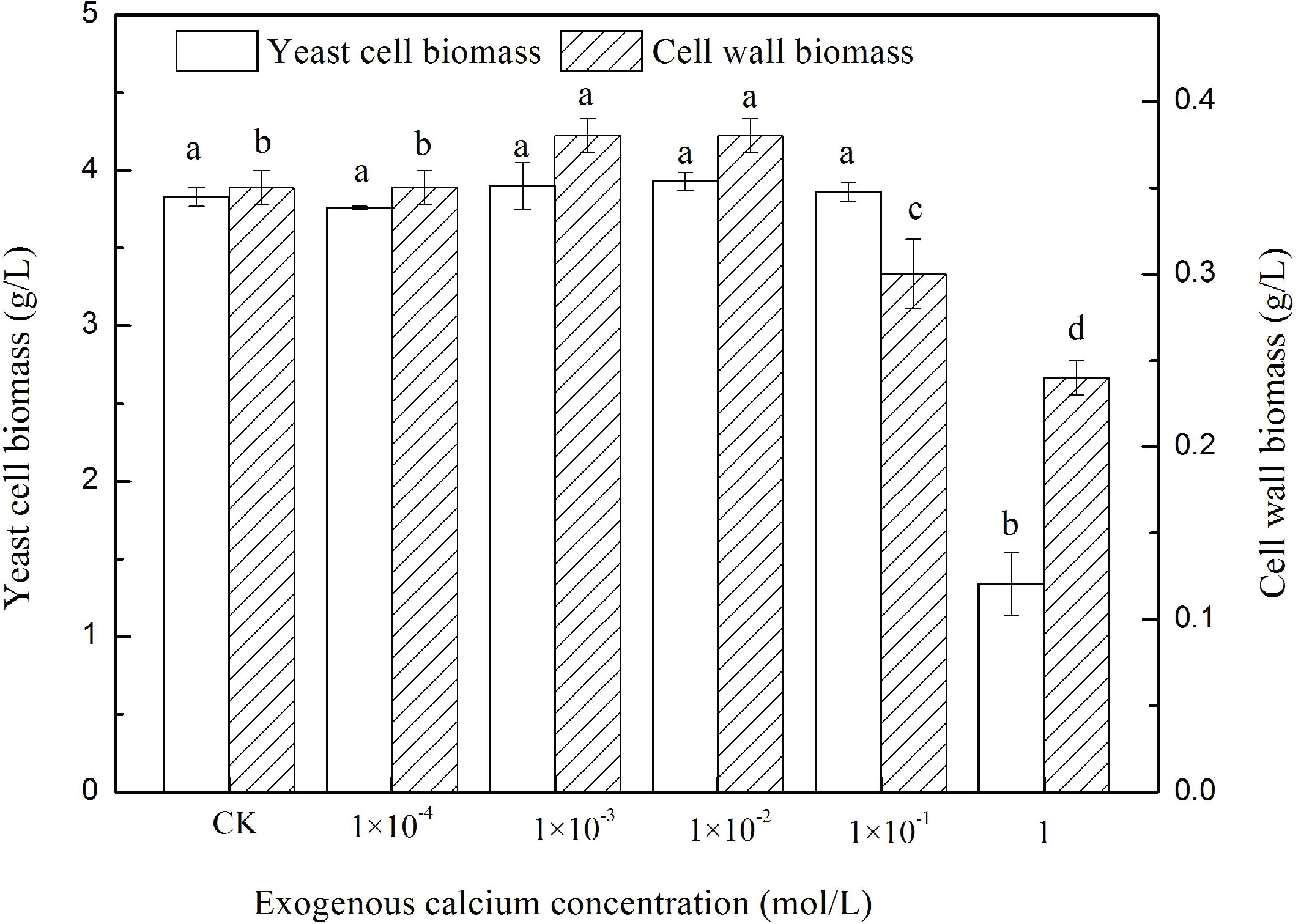
Different exogenous calcium culture cell biomass and cell wall biomass determination. Bars marked with different lowercase letters are significantly different (p < 0.05).

### 3.2 Intracellular calcium concentration determination

To confirm the effect of activation of exogenous calcium on the intracellular calcium pathway, the intracellular calcium concentration and the relation between intra- and extracellular calcium were determined. The fluorescence intensities of cytosolic free calcium in different exogenous calcium culture cells were determined at the same cell concentration of 1×10^5^/mL, and the results are shown in Figure 3. The relative intensity of cytosolic free calcium apparently intensified with increasing exogenous calcium concentrations to a certain extent, ranging from 1×10^-4^ to 1×10^-2^ mol/L. However, the fluorescence intensity of the cytosolic free calcium became weak and ultimately disappeared with the increase in the exogenous calcium concentration. The intracellular calcium had the highest concentration when the exogenous calcium concentration was 1×10^-2^ mol/L. This is due to the existence of a calcium steady-state system in *Saccharomyces cerevisiae*. During conditions of high exogenous calcium ion concentrations, the exogenous calcium ions could enter the yeast cells aided by transporter proteins ^[23, 24]^. However, under conditions of excessive intracellular calcium concentrations, the redundant calcium was partially transported into vacuoles using vacuolar proton pumps, and the other portion was transported extracellularly using Golgi/endoplasmic calcium pumps ^[25]^.

**Figure 3.**
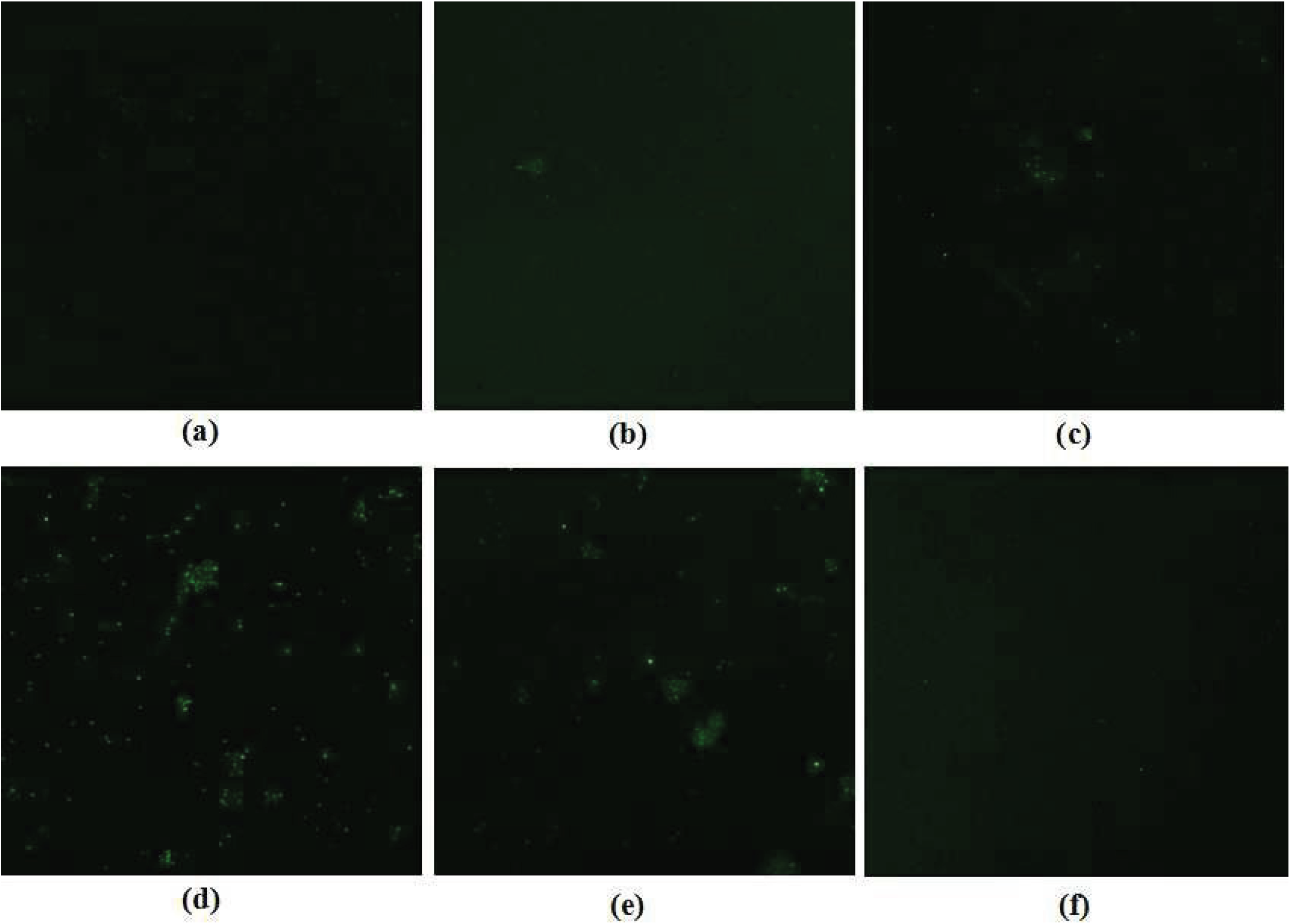
The fluorescence intensities of cytosolic free calcium in different exogenous calcium culture cells (a, b, c, d, e and f were 0, 1 × 10^-4^, 1 × 10^-3^, 1 × 10^-2^, 1 × 10^-1^, and 1mol/L, respectively).

### 3.3 The effect of exogenous calcium concentration on cell wall thickness

The ultrastructures of different exogenous calcium cultivated cells are shown in the TEM images at 25,000 × magnification (80.0 kv, 10.0 μA) in Figure 4, and the frequency histogram is used to calculate cell wall thickness (Figure 5). TEM images obviously displayed the tightness and thickness of the cell wall, and the cell wall thickness increased in parallel with the increasing calcium concentrations compared to the control group. However, the thickness reached its peak when the concentration of calcium added was 1×10^-2^ mol/L and then decreased as the calcium concentration continuously increased. In image *f* of the maximum calcium addition (1 mol/L), it appears that the cell wall layer was thinner and most easily damaged. The cell wall thickness values were determined by combining with the thickness statistics. The thickness values increased from 66.9 nm to 210.54 nm at 1×10^-2^ mol/L, and then decreased to 99.46 nm at the maximum calcium addition of 1 mol/L. These results indicated that the exogenous addition of calcium contributes to yeast cell wall layer formation to a certain degree because the increasing intracellular calcium could trigger the formation of the Ca^2+^-calcineurin complex in the cytoplasm, thus activating calcineurin ^[26]^. The activated calcineurin affected the formation of the cell wall by phosphorylating the transcription factor *crz1* and then regulating the expression of multiple downstream calcineurin-dependent genes ^[27, 28]^.

**Figure 4.**
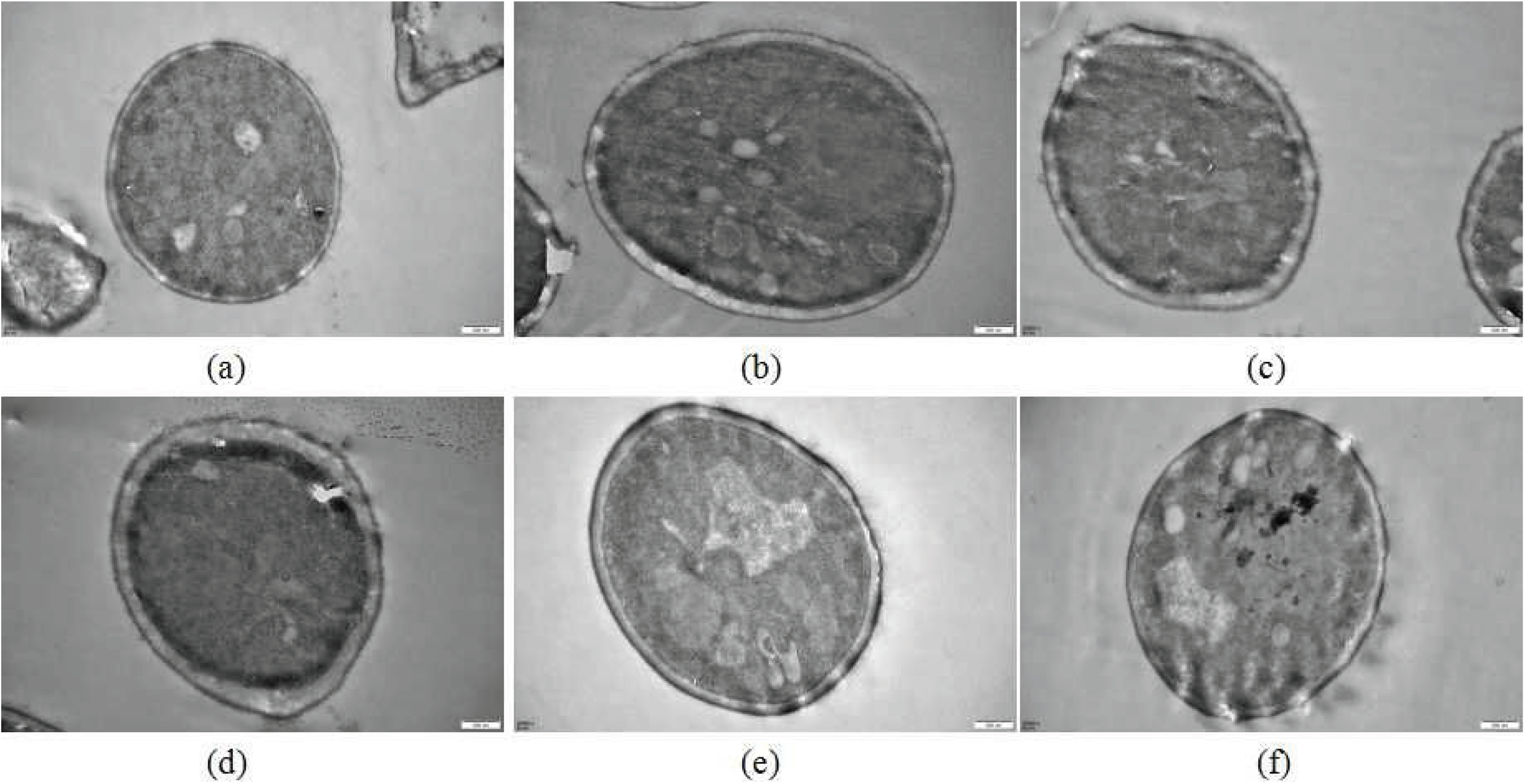
TEM images of different exogenous calcium culture cells at 25,000 × magnification (a, b, c, d, e and f were 0, 1×10^-4^, 1×10^-3^, 1×10^-2^, 1×10^-1^, and 1mol/L, respectively).

**Figure 5.**
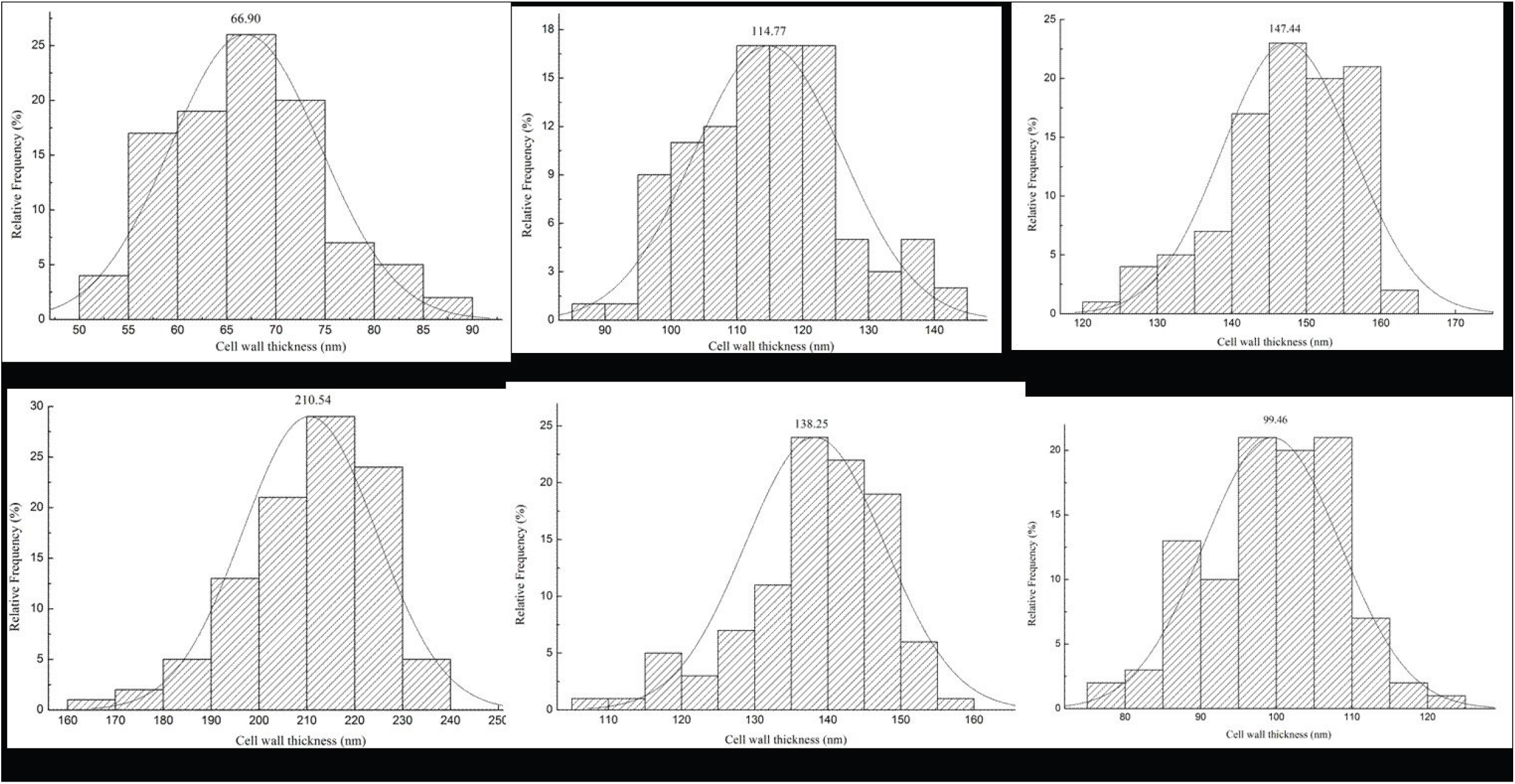
Histogram statistical results of different exogenous calcium culture cell wall thickness (a, b, c, d, e and f were 0, 1 × 10^-4^, 1 × 10^-3^, 1 × 10^-2^, 1 × 10^-1^, and 1mol/L, respectively).

### 3.4 The effect of exogenous calcium concentrations on the yeast cell wall insoluble network structure

Yeast cell walls are considered to be made up of different glucan types with different solubility properties. The solubility properties of yeast cell wall glucan had a direct relationship with the existence of chitin. Chitin, with its content less than 3%, connected with glucan by covalent bonds and could change soluble glucan to an insoluble state ^[29]^. Therefore, many studies have been conducted on the network structure composed of β-glucans (β-1,3-glucans and β-1,6-glucans) and chitin ^[18]^. Different exogenous calcium-cultivated cell wall network compositions were analyzed in this study, and the results of the β-1,3-glucan, β-1,6-glucan, and chitin contents are shown in Figure 6. As seen from the illustration, the β-glucan and chitin contents increased as the exogenous calcium concentration increased from 1×10^-4^ to 1×10^-2^ mol/L compared to the controls. Their contents then started to decrease at 1×10^-1^ mol/L, with a sudden drop at the calcium concentration of 1 mol/L. It displayed a similar trend with the results of the cell wall biomass and cell wall thickness since the insoluble β-glucan content influenced the density and thickness of the network structure and cell wall formation ^[11]^.

**Figure 6.**
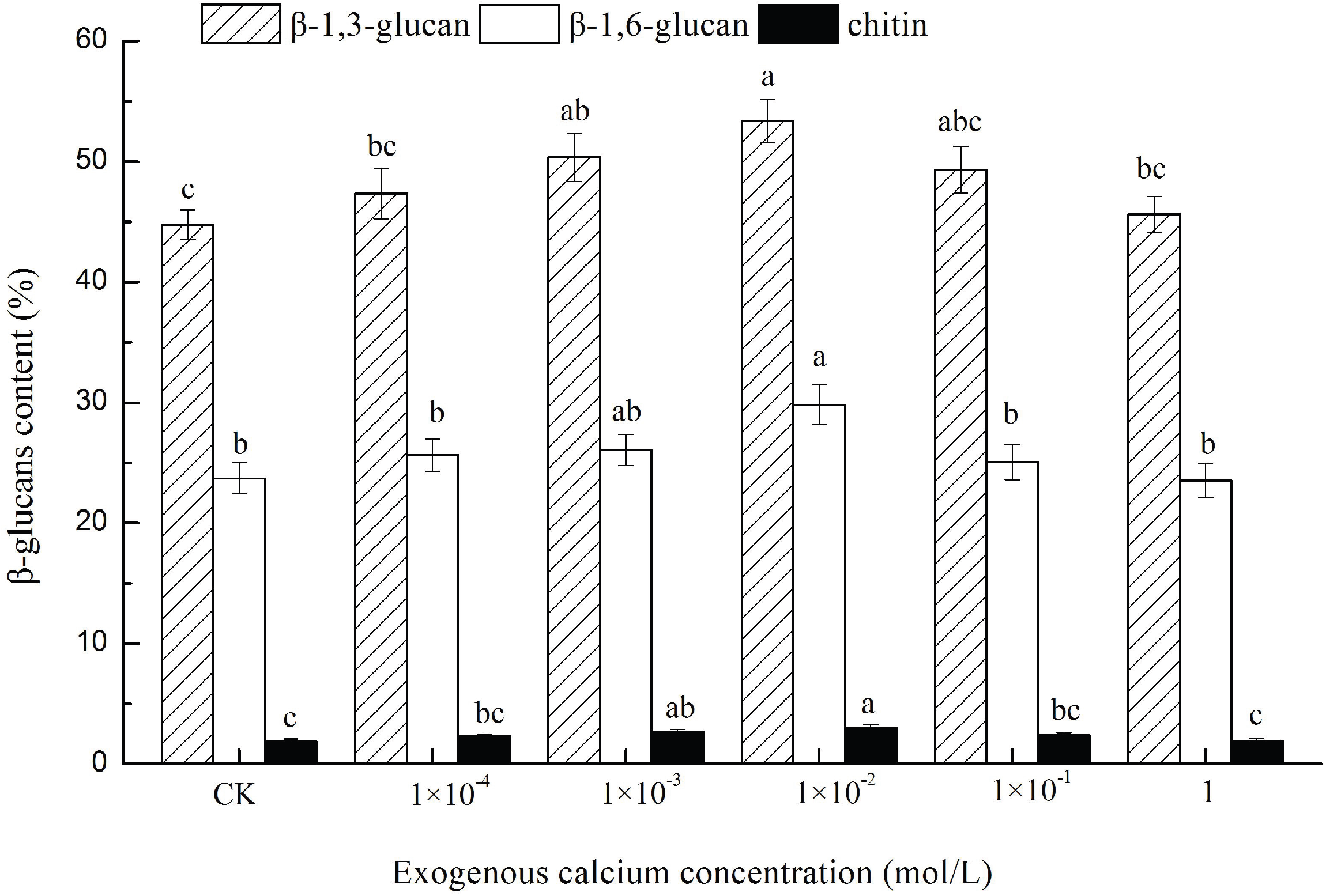
Different exogenous calcium culture cell wall insoluble network structure composition analysis. Bars marked with different lowercase letters are significantly different (p < 0.05).

### 3.5 The effect of exogenous calcium concentration on the activities of β-1,3-glucanase and β-1,3-glycosyl transferase

The cell wall network structure is biosynthesized with uracil diphosphate glucose as the only precursor substance and it then synthesizes long-chain soluble β-glucans utilizing β-1,3-glucanase, eventually forming insoluble β-glucans with chitin mediated by β-1,3-glycosyl transferase. Consequently, the activities of β-1,3-glucanase and β-1,3-glycosyl transferase play important roles during cell wall network structure formation ^[12]^. The effect of exogenous calcium concentrations on the β-1,3-glucanase and β-1,3-glycosyl transferase activities are shown in Figure 7. As shown in the figure, the activities of β-1,3-glucanase and β-1,3-glycosyl transferase increased as the exogenous calcium concentration increased within the concentration of 1×10^-2^ mol/L, and the results concurred with those observed in other studies ^[13]^. As the exogenous calcium concentration continued to increase (1×10^-1^ mol/L), the excessive calcium could act as osmotic pressure, which is harmful for yeast cells. The cell would be forced to stop its calcium response to promote the high osmotic pressure glycerol response (HOG) pathway since a high osmotic stress response attempts to repair the molecular damage and adapt to the new environment ^[30, 31]^. At this point, β-1,3-glucanase and β-1,3-glycosyl transferase, which are regulated by calcium signals, would suffer a sudden decrease, subsequently causing insoluble β-glucans and cell wall thickness and even a decrease in the cell wall and yeast biomass.

**Figure 7.**
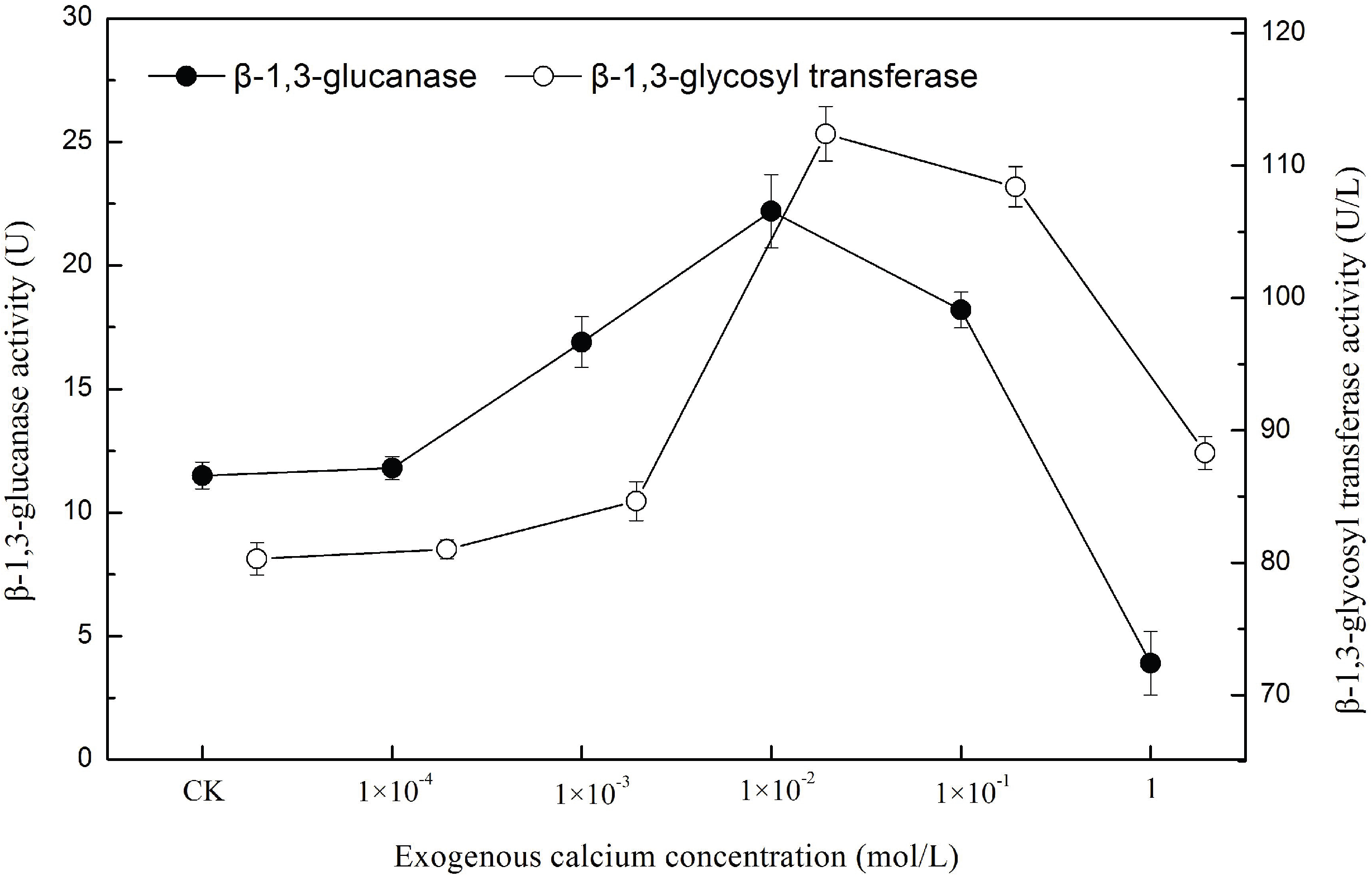
The activities of β-1,3-glucanase and β-1,3-glycosyl transferase in different exogenous calcium culture cells.

### 3.6 The effect of exogenous calcium concentration on patulin adsorption capability

The results of the patulin adsorption with different amounts of exogenous calcium-cultivated yeast cells are shown in Figure 8. All the cells tested could efficiently adsorb patulin from the aqueous solution and the apple juice, and the patulin adsorption ratios increased in parallel with the exogenous calcium concentration that increased within 1×10^-2^ mol/L. Patulin adsorption ratios increased from 83.4% to 94.8% in aqueous solution and from 76.2% to 85.9% in apple juice. Subsequently, the patulin adsorption ratios slightly decreased at 1×10^-1^ mol/L, with the adsorption ratios 92.5% and 83.7% in the aqueous solution and the apple juice, respectively. As the calcium ion concentration continued to increase to 1 mol/L, the patulin adsorption capability of the yeast cells dropped precipitously either in the aqueous solution or in the apple juice. It was evident that the effect of exogenous calcium on the patulin adsorption capability of the yeast cells was significant because the cell wall network structure and thickness changed with exogenous calcium concentration. The adsorption capability of the patulin increased in parallel with the density of the cell wall network structure ^[8]^. It can also be seen that the patulin adsorption capability of the yeast cells is greater in aqueous solutions. This is due to the nonspecific adsorption characteristic of yeast cells. In apple juice, a certain amount of pigments could be adsorbed as well, and they competed with the adsorption sites for patulin, thus removing the available adsorption sites in the yeast cell wall associated with patulin ^[6]^.

**Figure 8.**
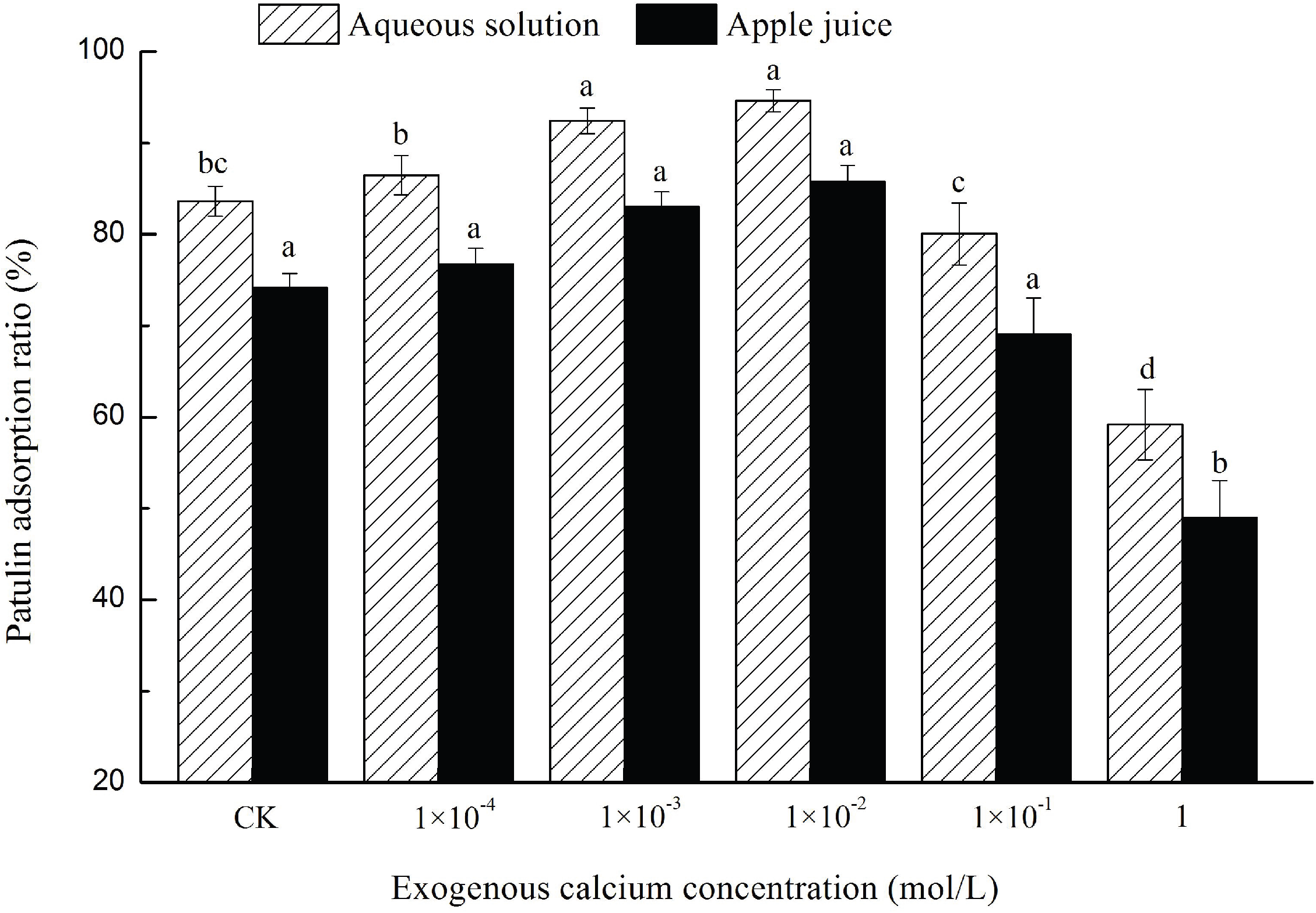
Patulin adsorption in aqueous solution and commercial apple juice by different exogenous calcium culture cells. Bars marked with different lowercase letters are significantly different (p < 0.05).

## 4. Conclusion

In this study, we have shown that exogenous calcium ions can improve the patulin adsorption capability of the yeast cell. This is the first report demonstrating a relationship between the calcium ion and patulin adsorption. Previous studies revealed that the patulin adsorption capability of yeast cells was primarily based on the cell wall filamentous network structure, and the adsorption process was considered to be due to the insertion of the free patulin adsorbed into the network pore structure ^[8]^. This peculiar network structure is formed with β-1,3-glucan and chitin, which were controlled by the activities of β-1,3-glucanase and β-1,3-glycosyl transferase ^[12]^. In the present study, research on the improvement of the ability to adsorb patulin was conducted by adding different concentrations of exogenous calcium ions during yeast growth. A series of experiments showed that the exogenous calcium ions could determine the insoluble network structure by regulating key enzyme activities under certain concentrations, thus indirectly influencing the yeast cell patulin adsorption capability.

We preliminarily speculated on the mechanisms of the improvement in the patulin adsorption capability from our results. An appropriate amount of exogenous calcium ions entered the yeast cell with the aid of transporter proteins on the cell membranes and subsequently activated the calcineurin pathway by combining with the target protein calcineurin ^[24]^. Furthermore, the activated calcineurin carried out multiple functions, including upregulating some genes that encode key enzymes associated with cell wall formation ^[16, 32]^. Nonetheless, superabundant exogenous calcium ions posed a threat to yeast cells since a high osmotic pressure glycerol response (HOG) pathway would be activated to respond to the superabundant calcium stress.

On the basis of this study, more questions raised require further study. For example, it is not clear whether exogenous calcium activates the calcineurin pathway to determine the patulin adsorption capability of the yeast cells. Additionally, it is unclear whether the decisive results obtained were mediated by a gene *crz1* regulatory effect. Intriguingly, one study showed that the β-1,3-glucanase regulatory gene *gsc2* and β-1,3-glycosyl transferase regulatory gene *crh1* were probably calcineurin dependent, and their activation and expression may be affected by the calcineurin/Crz1 pathway ^[22]^. Thus, determining if the key genes *gsc2* and *crh1* are upregulated with the concentration of exogenous calcium in *S. cerevisiae* would be of interest. Further studies to elucidate the calcineurin/Crz1 pathway and *crz1* gene expression driven by exogenous calcium are needed.

## Acknowledgments

This work was supported by “China Postdoctoral Science Foundation Funded Project” (2017M613055), “Postdoctoral Science Foundation Funded Project of Shaanxi Province” (2017BSHYDZZ46), and “The Fundamental Research Funds for the Central Universities” (GK201703066).

## Author Contributions

Conceptualization, Jianke Li; Methodology, Ying Luo; Software, Yanqing Han; Validation, Ying Luo, Xiaojiao Liu, Yanqing Han, and Jianke Li; Investigation, Ying Luo and Xiaojiao Liu; Data Curation, Ying Luo; Writing – Original Draft Preparation, Ying Luo; Writing – Review & Editing, Ying Luo and Xiaojiao Liu; Supervision, Jianke Li; Funding Acquisition, Ying Luo.

## References

1. Marin S, Mateo EM, Sanchis V, Valle-Algarr F M, Ramos A J, Jiménez M. 2011. Patulin contamination in fruit derivatives, including baby food, from the Spanish market. Food Chem 124: 563–568.

2. Wichmann G, Herbarth O, Lehmann I. 2002. The mycotoxins citrinin, gliotoxin, and patulin affect interferongamma rather than interleukin-4 production in human blood cells. Environ. Toxicol 17: 211–218.

3. Drusch S, Kopka S, Kaeding J. 2007. Stability of patulin in a juice-like aqueousmodel system in the presence of ascorbic acid. Food Chem 100(1): 192–197.

4. International Agency for Research on Cancer (IARC). 1993b. Monographs on the evaluation of carcinogenic risks of chemicals to humans. Lyon, France, 56: 489–521.

5. European Union. 2006. Commission regulation no. *1425/2003* amending commission regulation no. *1881/2006* setting maximum levels for certain contaminants in foodstuffs. Brussels Belgium L203: 1–3364.

6. Luo Y, Wang ZL, Yuan YH, Zhou ZK, Yue TL. 2015(a). Patulin adsorption of a superior microorganism strain with low flavor-affection of kiwi fruit juice. World Mycotoxin J 9(2): 195–203.

7. Var I, Erginkaya Z, Kabak B. 2009. Reduction of OTA levels in white wine by yeast treatments. J I Brewing 115: 30–34.

8. Luo Y, Wang JG, Liu B, Wang ZL, Yuan YH, Yue TL. 2015(b). Effect of yeast cell morphology, cell wall physical structure and chemical composition on patulin adsorption. Plos one 10(8): 1–16.

9. Yiannikouris A, Francois J, Poughon L, Dussap CG, Bertin G, Jeminet G, Pouany J. 2004(a). Adsorption of zearalenone by β-D-Glucans in the *Saccharomyces cerevisiae* cell wall. J Food Prot 67(6): 1195–1200.

10. Jouany JP, Yiannikouris A, Bertin G. 2005. The chemical bonds between mycotoxins and cell wall components of *Saccharomyces cerevisiae* have been identified. Archiva Zootechnica 8: 26–50.

11. Klis FM, Boorsma A, De Groot PW. 2006. Cell wall construction in *Saccharomyces cerevisiae*. Yeast 23: 185–202.

12. Mo HZ, Liu XY, Hu YJ. 2007. Advances in β-1,3-glucans biosynthesis of *Saccharomyces cerevisiae*i*s*. Food Sci 28(5): 358–362.

13. Li H. 2008. The purification, characteristic, and cDNA clone of β-1,3-glucanase in Thermophilus. Doctoral thesis. Shandong Agricultural University, Taian, China.

14. Cyert MS. 2003. Calcineurin signaling in *Saccharomyces cerevisiae*: how yeast go crazy in response to stress. Biochem Bioph Res Co 311: 1143–1150.

15. Miyakawa T, Mizunuma M. 2007. Physiological roles of calcineurin in *Saccharomyces cerevisiae* with special emphasis on its roles in G2/M cell-cycle regulation. Biosci Biotech Bioch 71: 633–645.

16. Ferreira RT, Courelas Silva, AR, Pimentel C, Batista-Nascimento L, Rodrigues-Pousada C, Menezes RA. 2012. Arsenic stress elicits cytosolic Ca^2+^ bursts and Crz1 activation in *Saccharomyces cerevisiae*. Microbiology 158: 2293–2302.

17. Manners DJ, Masson AJ, Patterson JC. 1973. The structure of a β-(1–3)-D-glucan from yeast cell walls. Biochem J 135: 19–30.

18. Yiannikouris A, Francois J, Poughon L, Dussap CG, Bertin G, Jeminet G, Pouany J. 2004(b). Alkali extraction of β-D-Glucans from *Saccharomyces cerevisiae* cell wall and study of their adsorptive properties toward zearalenone. J Agri Food Chem 52: 3666–3673.

19. MacDonald S, Long M, Gilbert J, Felgueiras I. 2000. Liquid chromatographic method for determination of patulin in clear and cloudy apple juices and apple puree: Collaborative study. J AOAC Int 83: 1387–1394.

20. Luo Y, Zhou ZK, Yue TL. 2017. Synthesis and characterization of nontoxic chitosan-coated Fe_3_O_4_ particles for patulin adsorption in a juice-pH simulation aqueous. Food Chem 221: 317–323.

21. Rita TF, Ana RC, Catarina P, Liliana BN, Claudina R, Regina AM. 2012. Arsenic stress elicits cytosolic Ca^2+^ bursts and Crz1 activation in *Saccharomyces cerevisiae*. Microbiology 158: 2293–2302.

22. Guillaume L, Howard B. 2006. Cell wall assembly in *Saccharomyces cerevisiae*. Microbio Mol Biol R 70 (2): 317–343.

23. Cui J, Kaandorpa JA, Ositelub O, Beaudry V, Knight A, Nanfacka YF, Cunningham KW. 2009. Simulating calcium influx and free calcium concentrations in yeast. Cell Calcium 45: 123–132.

24. Martin DC, Kim H, Mackin NA, Maldonado-Báez L, Evangelista CJ, Beaudry VG. 2011. New regulators of a high affinity Ca^2+^ influx system revealed through a genome-wide screen in yeast. J Biol Chem 286(12): 10744–10754.

25. Demaegd D, Foulquier F, Colinet AS, Gremillon L, Legrand D, Mariot P. 2013. Newly characterized Golgi-localized family of proteins is involved in calcium and pH homeostasis in yeast and human cells. P Natl Acad Sci USA 110(17): 6859–64.

26. Berchtold MW, Villalobo A. 2014. The many faces of calmodulin in cell proliferation, programmed cell death, autophagy, and cancer. Acta Bioch Bioph Sin 1843: 398–435.

27. Clapperton JA, Martin SR, Smerdon SJ, Gamblin SJ, Bayley PM. 2002. Structure of the complex of calmodulin with the target sequence of calmodulin-dependent protein kinase I: studies of the kinase activation mechanism. Biochem 41: 14669–14679.

28. Karababa M, Valentino E, Pardini G. 2006. Crz1, a target of the calcineurin pathway in *Candida albicans*. Mol Microbiol 59(5): 1429–1451.

29. Klis FM, Mol P, Hellingwerf K. 2002. Dynamic of cell wall structure in *Saccharomyces cerevisiae*. FEMS Microbiol Rev 26: 239–256.

30. Nissen, T. L., Hamann, C. W., Kielland-Brandt, M. C., Nielsen, J., Villadsen, J. Anaerobic and aerobic batch cultivations of *Saccharomyces cerevisiae* mutants impaired in glycerol synthesis. Yeast. 2000, 16 (5): 463–474.

31. Rebeca AM, Eliana R, Iwona W, Jan-Paul B, Willem HM, Marco S. 2001. Hyperosmotic stress response and regulation of cell wall integrity in *Saccharomyces cerevisiae* share common functional aspects. Mol Microbiol 41(3): 717–730.

32. Araki Y, Wu H, Kitagaki H, Akao T, Takagi H, Shimoi H. 2009. Ethanol stress stimulates the Ca^2+^-mediated calcineurin/Crz1 pathway in *Saccharomyces cerevisiae*. J Biosci Bioeng 107(1): 1–6.

